# Isolation, culture and maintenance of rabbit intestinal organoids, and organoid-derived cell monolayers

**DOI:** 10.1101/2020.07.15.205328

**Authors:** Egi Kardia, Michael Frese, Tanja Strive, Xi-Lei Zeng, Mary Estes, Robyn N. Hall

## Abstract

Organoids emulate many aspects of their parental tissue and have been used to study pathogen-host interactions, tissue development and regeneration, metabolic diseases, and other complex biological processes. Here, we report a robust protocol for the isolation, maintenance and differentiation of rabbit small intestinal organoids and organoid-derived cell monolayers. We also report conditions that sustain an intestinal stem cell population in spheroid culture. Rabbit intestinal spheroids and monolayer cultures propagated and expanded most efficiently in L-WRN-conditioned medium that contained the signalling factors Wnt, R-spondin and Noggin, and that had been supplemented with ROCK and TGF-β inhibitors. Organoid and monolayer differentiation was initiated by switching to a medium that contained less of the L-WRN-conditioned medium and was supplemented with ROCK and Notch signalling inhibitors. Using immunofluorescence staining and RT-qPCR, we demonstrate that organoids contained enterocytes, enteroendocrine cells, goblet cells and Paneth cells. These findings demonstrate that our rabbit intestinal organoids have many of the multi-cellular characteristics of, and closely resemble, an intestinal epithelium. This newly established organoid culture system will provide a useful tool to study rabbit gastrointestinal physiology and disease. For example, organoids and organoid-derived cells may be used to propagate and study caliciviruses and other enterotropic pathogens that cannot be grown in conventional cell culture systems.

## Introduction

The small intestine is lined by a simple, single-layer epithelium that is characterised by numerous finger-like protrusions (villi) and invaginations (crypts) that greatly enlarge the inner surface area of the small intestine (Fig 1). The differentiated epithelial cells of the villi have both absorptive and protective functions. The small intestinal epithelium contains four major types of specialised cells (enterocytes, enteroendocrine cells, goblet cells and Paneth cells), a pool of multipotent stem cells called crypt base columnar cells (CBCs) and a rare population of specialised epithelial cells that include tuft and M cells [1]. The abundance of each of these cell types varies within different segments of the small intestine. Enterocytes are the most abundant epithelial cell lineage in the small intestine and perform digestive and absorptive functions. The apical membrane of enterocytes is characterised by the presence of microvilli that form a ‘brush border’. This barrier is part of the host defence against luminal microbes and microbial toxins. Sucrase-isomaltase, lactase, maltase-glucoamylase and trehalase are some of the enzymes secreted from the apical surface of microvilli to aid nutrient absorption [2]. Enteroendocrine cells represent a small proportion of the cells in the epithelium of the small intestine. Like enterocytes, they are tall and columnar in appearance with a microvilli-covered apical surface that stands in direct contact with the intestinal lumen. In contrast to enterocytes, on the basolateral side, enteroendocrine cells are equipped with a chemosensory extension called a neuropod that extends toward the enteric nervous system in the underlying mucosal layer [3]. Enteroendocrine cells have secretory granules that store peptides such as chromogranin A [4] and/or hormones that are released into the blood stream in response to food intake [5]. Goblet cells are mucin-producing cells found scattered in the small intestinal epithelium lining. A goblet cell has a narrow base and an oval apical portion that contains mucin granules. Mucus secreted from these cells forms a gel-like coating over the surface epithelium that protects against pathogen invasion [6]. Mucin 2 (Muc2) and Muc5ac are the major components of the mucus in the gastrointestinal tissue [7]. Paneth cells reside at the base of the crypts and provide survival signals to adjacent crypt stem cells. Paneth cells also play a role in the innate immune defence; the cells’ secretory granules contain several anti-microbial agents, including α-defensins and lysozyme [8]. These host defence proteins are pro-inflammatory mediators that help to protect the host against enteric pathogens. CBCs, the intestinal stem cell population, are usually found at the crypt base, intermingled with Paneth cells. CBCs regenerate the small intestinal epithelium through self-renewal and differentiation into specialised epithelial cells. The generation of new epithelial cells near the base of a crypt pushes older cells up towards the tip of the neighbouring villi, a process that continuously replaces senescent cells, which are lost to apoptosis [9,10]. Lgr5 and CD44 are two of the most common stem cell markers that can be used to identify small intestinal CBCs. Lgr5 was at first shown to be expressed in actively cycling CBCs [11], however later become a global marker of adult stem cells since they can also be found in the proliferative compartment of multiple other organs [12]. In addition, CD44 is also highly expressed in both mouse and human stem cell populations of the small intestine [13].

**Fig 1.**
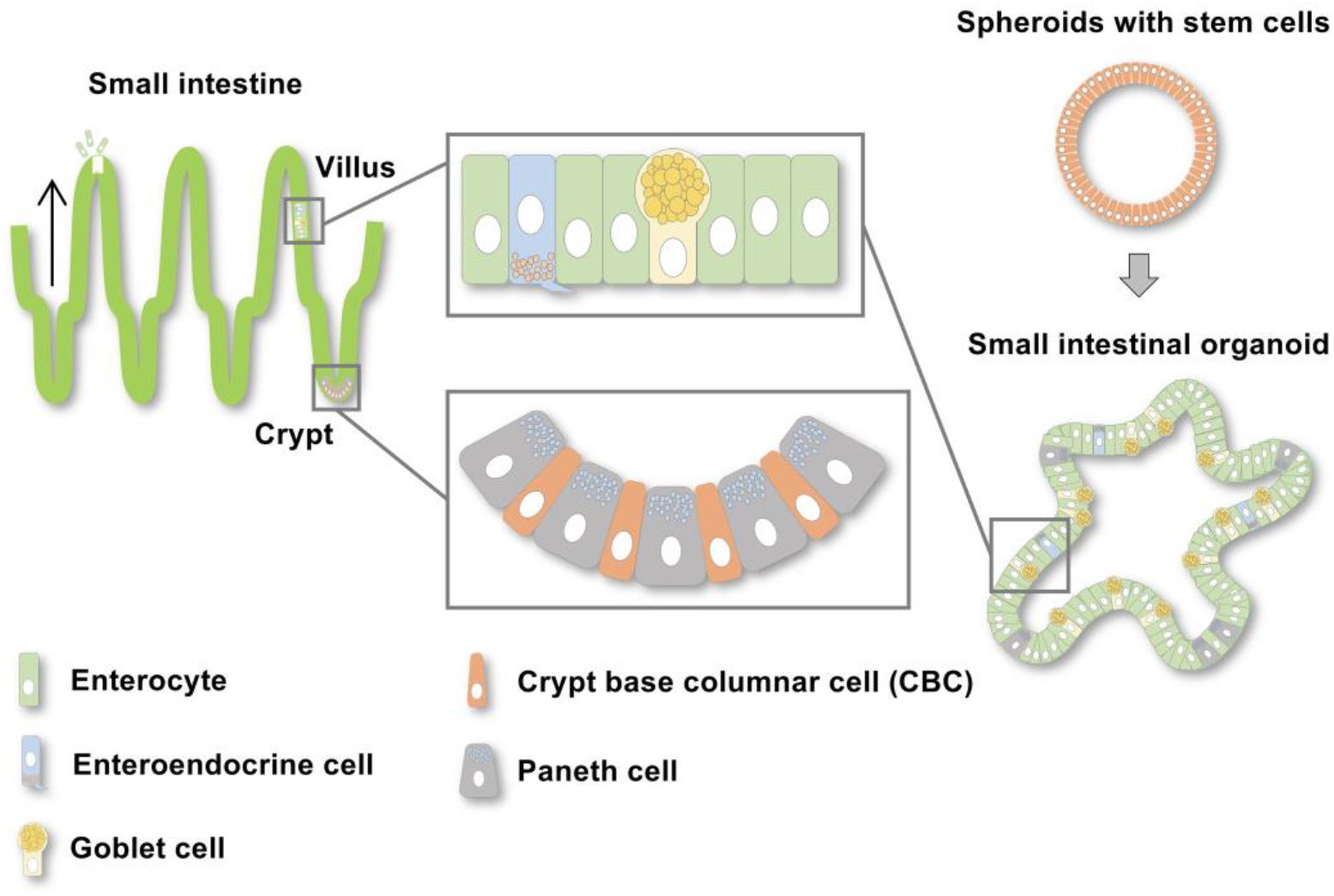
Epithelial cell types in the small intestine. The epithelial layer of the small intestine is organised into villi and crypts. Villi consist of differentiated epithelial cells including enterocytes, enteroendocrine cells and goblet cells. The crypts contain Paneth cells and crypt base columnar cells (CBCs). The CBCs continuously proliferate and provide new cells that move up the neighbouring villi during differentiation (black arrows). Older cells are pushed to the tip of the villi from where cells are continuously shed into the lumen to make room for the next generation of epithelial cells. Organoids generated from small intestinal stem cells should ideally contain all types of epithelial cells present in the intestine, including enterocytes, enteroendocrine cells, goblet cells and Paneth cells.

Two-dimensional (2D) cell culture models suffer many disadvantages, including the lack of a tissue-specific architecture. They typically lack an extracellular matrix (ECM) and many of the cell–cell interactions that occur *in vivo*. Organoids are artificial three-dimensional (3D) tissue constructs that can be generated from self-organising stem cells in culture and that allow the generation of near-physiological conditions in cell culture [14]. This has allowed researchers to conduct both basic and translational research that was previously thought to be impossible. Organoids present significant prospects for modelling tissue physiology and pathology, microbial infections, toxicology studies and drug discovery. In many instances, organoids already have, or will, replace animal models that are expensive, laborious and have considerable animal welfare implications. Differentiated mature organoids (herein called organoids) contain all major cell types of the tissues from which the stem cells were isolated. Organoids can be generated from pluripotent stem cells that are either embryo-derived or isolated from adult stem cells retrieved from tissue biopsies [14,15]. When supplemented with appropriate growth factors and cultured in an ECM, these stem cells can self-renew and build a tissue-like structure that recapitulates many features of their original environment. Intestinal organoids, for example, contain many of the differentiated epithelial cells known from adult intestine tissues, including enterocytes, goblet cells, enteroendocrine and Paneth cells (Fig 1).

In mice intestinal organoids, CBCs must first undergo self-organisation into symmetrical sphere-like structures, herein referred to as spheroids. Then, Paneth cells arise from CBCs; Paneth cells guide the differentiation of other cells and drive the formation of multilobular structures that resemble the crypt-and-villus architecture of the intestine [16,17]. The tissue-specific microenvironment is a key factor that drives stem cell differentiation *in vivo*. To grow organoids, a suitable environment must be created, which can be achieved by using an artificial ECM and a combination of tissue-specific growth factors [18]. Most commercially available ECMs contain a heterogeneous mixture of matrix proteins (e.g., laminin, collagen IV, entactin and proteoglycans) and growth factors that are harvested from cultured Engelbreth-Holm-Swarm mouse sarcoma cells. In human intestinal organoids, critical signalling pathways depend on Wnt, R-spondin and Noggin [19], with Wnt being crucially important for maintaining the proliferation of a healthy stem cell pool [20,21]. Wnt signalling is highly conserved across metazoans, regulating embryonic development and adult tissue homeostasis. The canonical Wnt or Wnt/β-catenin pathway is activated when Wnt proteins bind to the Frizzled receptor family [22,23]. This signal can be further enhanced through the binding of R-spondin proteins [23], and this signalling enhancement is required to drive the differentiation of stem cells in cell culture [24]. Noggin is another growth factor that is needed to maintain an intestinal stem cell pool [20]. Noggin interferes with the binding of bone morphogenetic proteins (BMPs) to their receptor [25], thus antagonising the function of cytokines that restrict intestinal stem cell proliferation [20,26]. The removal of Wnt, R-spondin and Noggin signalling allows stem cells to develop from an undifferentiated to a differentiated state, and thereby triggers the formation of organoids from spheroids.

Here, we report the development of robust protocols for the isolation, maintenance and long-term cryogenic storage of rabbit small intestinal spheroids from duodenum, jejunum and ileum segments, the differentiation of duodenal spheroids to organoids, and the cultivation of organoid-derived cell monolayers.

## Materials and methods

### Ethics statement

This study was approved by the CSIRO Wildlife and Large Animal Ethics Committee (permit numbers #2016-22 and #2016-02). All animal procedures were carried out at CSIRO Black Mountain Laboratories according to the Australian Code for the Care and Use of Animals for Scientific Purposes.

### Animals

A total of three adult European rabbits (*Oryctolagus cuniculus*) were used for this study: one “New Zealand white” laboratory rabbit (male, 3.63 kg) and two wild rabbits (one female, 1.9 kg and one male, 1.5 kg). The laboratory rabbit was euthanised by intravenous injection of Lethabarb™ (Virbac, Carros, France) following intramuscular anaesthesia with 30 mg/kg ketamine (Mavlab, Queensland, Australia) and 5 mg/kg xylazine (Troy Laboratories, New South Wales, Australia). Wild rabbits were opportunistically sampled during routine control operations (rabbit shooting) in a nearby National Park.

### Isolation and cultivation of intestinal epithelial cells

The isolation of intestinal epithelial cells was performed as described by Miyoshi and Stappenbeck, 2013 [27] with modifications (Fig 2). Briefly, the duodenum, jejunum and ileum were collected from a laboratory rabbit and dissected using sterile surgical scissors and tissue forceps. Tissue samples were placed in a 50-ml tube containing ice-cold sterile phosphate buffered saline (PBS) supplemented with 100 μl/ml antibiotic/antimycotic solution containing 10,000 units/ml of penicillin, 10 mg/ml of streptomycin and 25 μg/ml amphotericin B (Sigma-Aldrich, Missouri, USA). The excess fat surrounding the tissue was removed and the intestinal lumen was flushed with ice-cold PBS using a 10-ml syringe with an 18-G blunt needle. The cleaned intestine samples were then opened lengthwise, cut into 1 × 1 cm pieces and incubated overnight in digestion medium containing 1 mg/ml collagenase type I (Gibco, Massachusetts, USA) and 100 μl/ml antibiotic/antimycotic solution in Modified Hank’s Balanced Salt Solution (HBSS) (Sigma-Aldrich). Duodenum samples from wild rabbits were processed at the collection site in a similar manner and transported to the laboratory in a cooler box with ice packs (jejunum and ileum were not collected from wild rabbits). Tissue pieces were digested at 4 °C on an orbital shaker-incubator at 200 *×* rpm (Bioline Global, New South Wales, Australia). After overnight digestion, a cell scraper was used to dislodge the epithelium from the intestine. The epithelial cells were transferred into a 50-ml tube containing 0.25% trypsin/EDTA (Gibco), incubated for 5 min at 37 °C and passed through a 70-μm cell strainer. The digestion was stopped by adding 10% foetal bovine serum (FBS; Gibco) and the cells were pelleted by centrifugation at approximately 250 × g for 5 min at 4 °C (Eppendorf 5804 R, Hamburg, Germany). After resuspension, red blood cells were removed using ammonium chloride (red blood cell lysing buffer; Sigma-Aldrich). The cells were pelleted again by centrifugation at approximately 250 × g for 5 min at 4 °C. The epithelial cell pellets were then washed twice with PBS, centrifuged at approximately 250 × g for 5 min at 4 °C and resuspended in thawed Geltrex™ LDEV ([lactose dehydrogenase elevating virus]-Free Reduced Growth Factor Basement Membrane Matrix; Gibco). Two 15-μl drops of the matrix-cell suspension were pipetted into wells of a Nunc™ 24-well-Nuclon Delta-treated plate (Thermo Fisher Scientific, Massachusetts, USA) and allowed to solidify for 15 min at 37 °C before 400 μl of proliferation medium (described in the next section) was added to each well. The cultures were monitored daily to assess the formation of spheroids. The proliferation medium was changed every 3 days until the matrix dome became crowded with spheroids.

**Fig 2.**
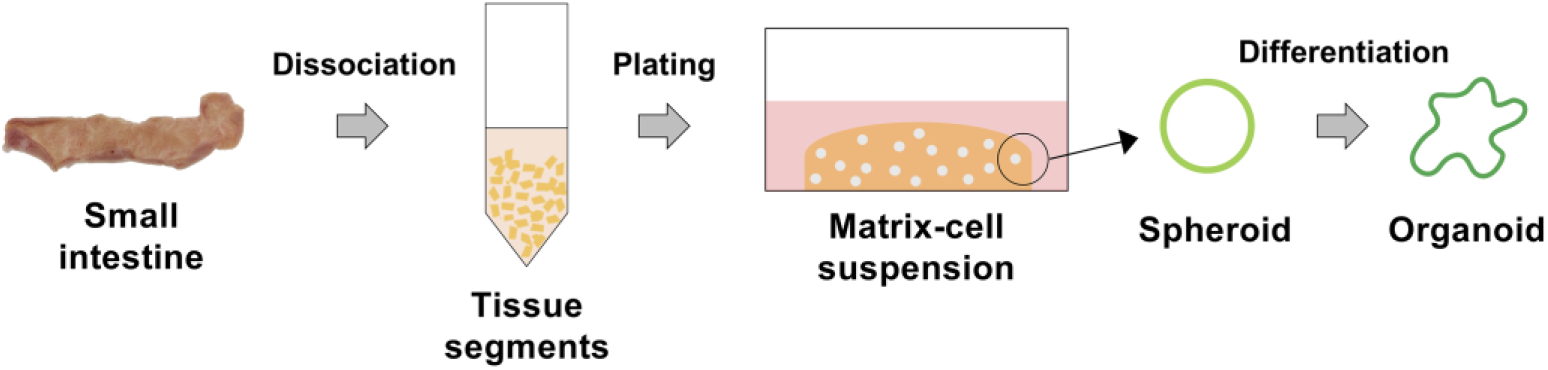
Generation of rabbit small intestinal organoids. Sections of the small intestine were cut and incubated in digestion medium overnight. The dissociated intestinal epithelial cells were cultured in ECM. L-WRN-conditioned medium was added to initiate spheroid formation. Differentiation medium was used to induce the formation of mature differentiated intestinal organoids.

### Proliferation medium

A conditioned medium was produced using a mouse fibroblast cell line that was genetically modified to express and secrete Wnt3a, R-spondin 3 and Noggin (abbreviated L-WRN; ATCC^®^ CRL-3276™, Virginia, USA) [27]. The proliferation medium was prepared by diluting the conditioned medium 1:1 with basal medium that contained Advanced DMEM/F12 (Gibco), 20% FBS (v/v) and 2 mM GlutaMAX™ Supplement (Gibco). Spheroids of the small intestine were cultured and expanded in conditioned medium supplemented with 10 μM of Rho-associated protein kinase (ROCK) inhibitor (Y-27632; Cayman Chemicals, Michigan, USA) and 10 μM of transforming growth factor-β (TGF-β) type I receptor inhibitor (SB-431542; Cayman Chemicals).

### Passaging and cryopreservation of confluent intestinal spheroid cultures

Spheroid cultures were split and sub-cultured with fresh proliferation medium every week, or sooner if dead cells started to accumulate in the lumen. Briefly, the old medium was removed without disturbing the matrix dome and the dome was carefully washed using basal medium. To dissolve the matrix and dissociate the spheroids, TrypLE™ Express Enzyme (Gibco) was added and the dome was broken up by gently pipetting the enzyme solution up and down until the spheroids were released from the matrix. The spheroids were transferred into a new 15-ml tube and incubated for 10–15 min at 37 °C. The digestion was stopped by adding 10% FBS and the cells were pelleted by centrifugation at about 100 *×* g for 5 min at 4 °C. For passaging, the cell pellets were resuspended in thawed Geltrex™ and the matrix-cell suspension was transferred into a 24-well-plate (two 15-μl drops containing 1 × 10^5^ cells per well). The matrix was allowed to solidify for 15 min at 37 °C before 400 μl of fresh proliferation medium was added. To freeze dissociated spheroids, cell pellets were resuspended in Recovery™ Cell Culture Freezing Medium (Gibco) and 1 × 10^6^ cells were transferred to a cryovial. The cryovial was then placed in a freezing container at −80 °C overnight and transferred to liquid nitrogen for long-term storage.

### Differentiation medium

To promote the maturation and differentiation of duodenal spheroids to organoids, proliferation media was replaced by differentiation media that contained less L-WRN-conditioned medium (i.e., 5%, compared to 50% in proliferation media) and that was supplemented with 10 μM of ROCK inhibitor and 50 ng/ml of DAPT (Notch signalling inhibitor; Cayman Chemicals). Although we initially established spheroid cultures from a laboratory rabbit (duodenum, jejunum, and ileum tissues) and two wild rabbits (duodenum tissues), we subsequently focussed our study on the characterisation of duodenal spheroids and organoids from the laboratory rabbit. Duodenal spheroids were incubated in differentiation medium for 4 days to establish differentiated and mature organoids.

### Transforming duodenal spheroids to monolayer cultures

Monolayers of spheroid-derived cells were cultured in either flat-bottom, tissue culture-treated 96-well plates (Corning, New York, USA) or in 6.5-mm Corning^®^ Transwell^®^ polycarbonate membrane cell culture inserts placed in conventional 24-well-plates (Corning). Prior to use, both plates and inserts were coated with 100 μg/ml collagen type IV (Sigma-Aldrich) for at least 2 h at 37 °C. To generate cells for monolayer cultures, proliferating spheroids were grown for 4–7 days and digested using TrypLE™ Express Enzyme, as described above. The resulting crude cell suspension was passed through a 70-μm cell strainer into a 50-ml tube and dissociated cells were counted using a haematocytometer. Approximately 7 × 10^4^ cells in a 100-μl drop of proliferation medium were placed either in the middle of a well of a 96-well plate or in a Transwell^®^ membrane insert. For Transwell^®^ cultures, an additional 600 μl of proliferation medium was added to the lower compartment of the plate. Differentiation of the cells was initiated by changing the proliferation medium to differentiation medium after the cells reached confluency (typically within 1 or 2 days). As expected, the media change triggered cell differentiation. Most cells differentiated within 3 days and the number of differentiated cells increased further with time; however, cells started to die 5 days after the switch to differentiation medium.

### Histological analysis

At selected time points (day 4–7 for spheroids and day 4 for organoids), spheroids and mature organoids were fixed and stained for histological examination (Fig 3). Briefly, the medium was discarded, the matrix dome containing the organoids was washed with PBS and the matrix was gently broken up by pipetting through 1,000-μl pipette tips with their end removed to increase the bore size. The organoids were then transferred to a 15-ml tube, pelleted by centrifugation at approximately 100 × g for 5 min at 4 °C, fixed in 4% paraformaldehyde for 2 h at 4 °C and embedded in PBS containing 3% low-melting point agarose (Bio-Rad, California, USA). The agarose gels containing the fixed organoids were processed in a standard automated tissue histology processor, embedded in paraffin and sectioned into 3-μm thick slices using a semi-automated rotary microtome (Leica Biosystems, Wetzlar, Germany). The slices were floated in a 37 °C water bath and mounted either on frosted-edge slides for histological staining (Hurst Scientific, Forrestdale, Western Australia) or on poly-L-lysine-coated slides for immunostaining (Thermo Fisher Scientific). The mounted sections were dried overnight at room temperature.

**Fig 3.**
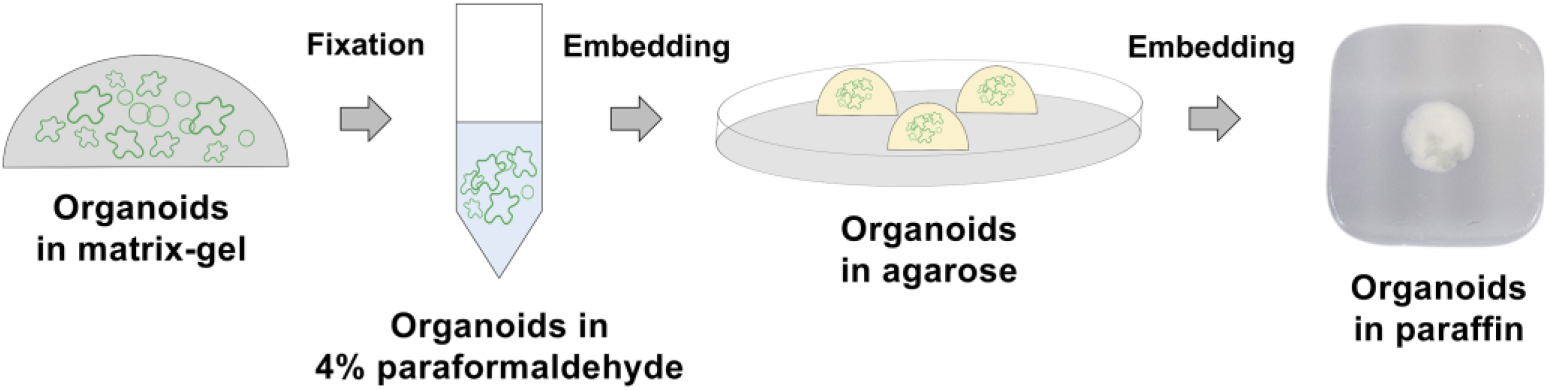
Histology and immunofluorescence staining. Organoids were fixed in 4% paraformaldehyde for 2 h before embedding in 3% agarose and then paraffin.

### Haematoxylin and eosin (H&E) staining

H&E staining was performed to evaluate the structure and composition of the spheroids and organoids. The procedure involved a 2-min deparaffinisation step with xylene (Sigma-Aldrich), followed by a rehydration step with graded ethanol solutions (100%, 90%, 80% and 70%) and tap water (each for 2 min). Next, samples were stained with Harris’ haematoxylin (Sigma-Aldrich) for 5 min, followed by incubation with 0.5% acid alcohol for 2 sec, tap water for 2 min, 0.2% ammonia water (Sigma-Aldrich) for 20 sec and again tap water for 2 min. Samples were counter-stained with eosin (Sigma-Aldrich) for 3 min and dehydrated using graded ethanol solutions (70%, 80%, 90% and 100%; each for 10 sec), followed by a final clearing step with xylene for 2 min, before the slides were mounted using dibutylphthalate polystyrene xylene (DPX; Sigma-Aldrich).

### Periodic acid-Schiff (PAS) staining

PAS staining was performed to identify the mucus-producing cells in organoids and differentiated monolayers grown in the 4-well Nunc^®^ Lab-Tek^®^ II Chamber Slide™ system (Thermo Fisher Scientific). The procedure involved a deparaffinisation step with xylene for 2 min (this step was skipped for monolayer staining), followed by a rehydration step with graded ethanol solutions (100%, 90%, 80% and 70%) and tap water (each for 2 min). Next, samples were incubated with 0.5% periodic acid (Sigma-Aldrich) for 5 min, tap water for 1 min, Schiff’s reagent (Sigma-Aldrich) for 15 min and again tap water for 5 min. Samples were counter-stained with Harris’ haematoxylin for 30 sec and dehydrated using graded ethanol solutions (70%, 80%, 90% and 100%; each for 10 sec). Lastly, the samples were cleared using xylene for 2 min and mounted using DPX.

### Immunofluorescence

Monolayers grown in collagen type IV-coated chamber slides were washed twice with PBS and incubated in 4% paraformaldehyde in PBS for 15 min at room temperature, then washed again twice with PBS before immunostaining. Spheroid/organoid sections on poly-L-lysine-coated glass slides were heated to 60 °C and dipped in xylene for 3 min to remove the surrounding paraffin. The sections were rehydrated with graded ethanol solutions (100%, 90%, 80% and 70%) and PBS (each for 3 min). After paraffin removal and rehydration, both spheroid/organoid sections and monolayers in chamber slides were placed in 10 mM citrate buffer for antigen retrieval and incubated for 20 min at 95 °C to expose antigenic sites. Following antigen retrieval, the slides were allowed to cool, then washed with PBS for 5 min. The slides were then permeabilised with 0.25% Triton X-100 (Sigma-Aldrich) in PBS for 10 min at room temperature, washed with PBS for 5 min and blocked with 5% bovine serum albumin (BSA; Sigma-Aldrich) in PBS-Tween 20 (Sigma-Aldrich) for 20 min. The following antibodies were used: anti-E-cadherin (5 μg/ml; LS-Bio, Washington, USA), anti-CD44 (10 μg/ml; Thermo Fisher Scientific), anti-mucin 5ac (5 μg/ml; Abcam, Cambridge, United Kingdom), anti-sucrase-isomaltase (10 μg/ml; Abcam) and anti-chromogranin A (25 μg/ml; Thermo Fisher Scientific); for further details, see Table 1. All primary antibodies were diluted in PBS-Tween 20 containing 1% BSA and placed into a humidified chamber and incubated overnight at 4 °C. Slides were then washed two times with PBS for 5 min, and then appropriate secondary antibodies, diluted in PBS with 1% BSA, were added. The slides were incubated at room temperature for another 40 min, washed two times with PBS, counterstained with 4’,6-diamidino-2-phenylindole (DAPI) (Sigma-Aldrich) for 5 min at room temperature and mounted with Fluoroshield™ (Sigma-Aldrich).

**Table 1.** Antibodies used for intestinal organoid characterisation.

| Antibody name | Cell specificity | Product number | Supplier | Conjugated |
| --- | --- | --- | --- | --- |
| Rabbit polyclonal anti-CDH1/E cadherin | Surface antigen of epithelial tissues, including the GI tract | LS-C351977 | LS Bio | Unconjugated |
| Rat monoclonal anti-CD44 | Surface antigen of stem cells | MA4400 | Thermo Fisher Scientific | Unconjugated |
| Rabbit polyclonal anti-sucrase-isomaltase | Enterocyte brush border enzyme | ab98872 | Abcam | Unconjugated |
| Mouse monoclonal anti-chromogranin A | Secretory vesicles of endocrine cells | MA5-13096 | Thermo Fisher Scientific | Unconjugated |
| Mouse monoclonal anti-mucin 5AC | Mucus produced in goblet cells | ab212636 | Abcam | Unconjugated |
| Secondary goat anti-rabbit | Goat polyclonal secondary antibody to rabbit IgG | ab150077 | Abcam | Alexa Fluor 488 |
| Secondary goat anti-rabbit | Goat polyclonal secondary antibody to rabbit IgG | ab150078 | Abcam | Alexa Fluor 555 |
| Secondary goat anti-rat | Goat polyclonal secondary antibody to rat IgG | A-11006 | Thermo Fisher Scientific | Alexa Fluor 488 |
| Secondary goat anti-rat | Goat polyclonal secondary antibody to rat IgG | A-21434 | Thermo Fisher Scientific | Alexa Fluor 555 |
| Secondary goat anti-mouse | Goat polyclonal secondary antibody to mouse IgG | ab150113 | Abcam | Alexa Fluor 488 |
| Secondary goat anti-mouse | Goat polyclonal secondary antibody to | A-21424 | Thermo Fisher | Alexa Fluor 555 |
|  | mouse IgG |  | Scientific |  |

### Imaging analysis

All staining was examined under a bright field phase-contrast/fluorescence inverted TI-U microscope (Nikon, Tokyo, Japan). Image analysis and processing was performed with NIS-Element software (Nikon).

### RT-qPCR

Spheroids and organoids were harvested using TrypLE™ and centrifuged at approximately 250 × g for 5 min at 4 °C to form a pellet. Total RNA was isolated using the RNeasy Mini kit (Qiagen, Hilden, Germany), which included a genomic DNA digestion step with RNase-free DNase (Qiagen), as per the manufacturer’s instructions. Primers for RT-qPCR were designed to be intron spanning and between 17–21 bases in length using NCBI Primer-BLAST (primer sequence information is provided in Table 2). RT-qPCRs were carried out using the SensiFAST SYBR No-ROX kit according to the manufacturer’s instructions (Bioline, London, United Kingdom) on a CFX96™ real-time instrument (Bio-Rad). The following cycling conditions were used: denaturation at 95° C for 10 sec min, amplification at 63° C for 40 sec and extension at 78° C for 10 sec (40 cycles). ‘No template’ and ‘no reverse transcriptase’ controls were performed with each run. Relative gene expression analysis was performed with three biological and three technical replicates for each experimental condition (i.e., proliferating spheroids, proliferating monolayers, differentiated organoids and differentiated monolayers) and was calculated using the 2^-ΔΔCt^ method using CFX Maestro software (Bio-Rad). Transcription was normalised to the expression levels of the housekeeping gene (18S ribosomal RNA) in proliferating spheroids/monolayers.

**Table 2.**
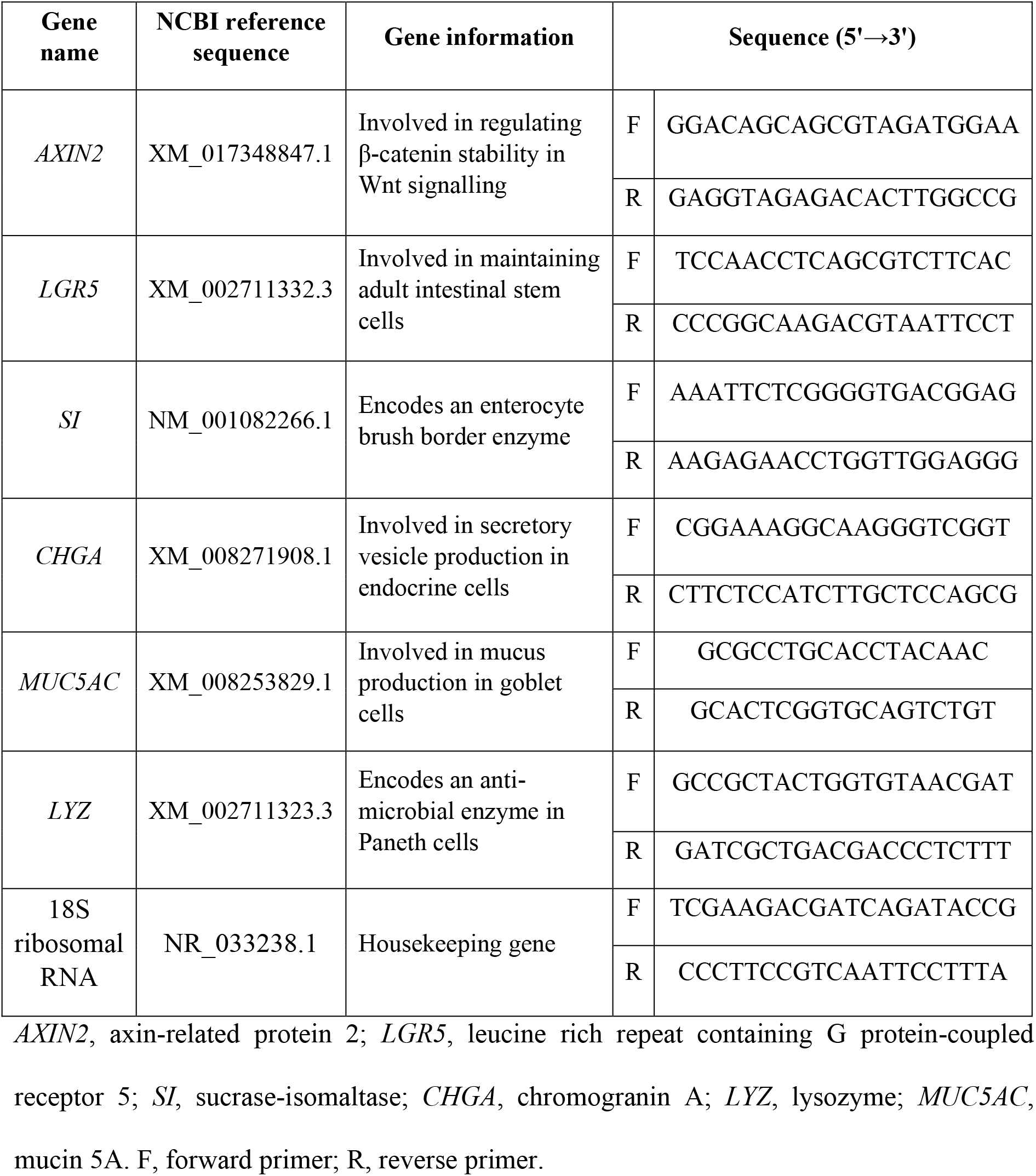
Primer sequences used for rabbit gene expression analyses.

### Statistical analysis

The diameter of spheroids in different culture conditions was measured using NIS-Element software (Nikon). The average diameter of rabbit intestinal spheroids was calculated from fifty representative examples per well for three biological replicates and statistical significance was analysed using Student’s t-test (significance was defined as *p* < 0.05). Graphs were generated and the statistical analyses were performed using GraphPad Prism Version 8.0 (GraphPad software, California, USA).

## Results

### Rabbit intestinal spheroid morphology

Intestinal organoid models derived from ‘exotic’ animals such as such as pigs [28], horses [29,30], cats, dogs, chicken [30] and ferrets [31] have been described previously. These were generated to recreate the species-specific molecular and histological phenotypes seen *in vivo*. Here we report a protocol for the generation and cultivation of 3D rabbit intestinal spheroids and organoids. Laboratory and wild rabbits were used to isolate small intestinal epithelial cells. When these cells were cultured in ECM in the presence of WRN factors, spheroids started to form within 2 days. We generated intestinal spheroids from duodenum, jejunum and ileum tissue segments from a laboratory rabbit (Fig 4A–C) and duodenal spheroids from wild rabbits (Fig 4D). These spheroids consisted of a monolayer of epithelial cells surrounding a liquid-filled lumen (Fig 5A, B), as demonstrated by E-cadherin expression (Fig 5C, D). In addition, these spheroids also contained a highly proliferative CD44^+^ stem and/or progenitor cell population (Fig 5E, F).

**Fig 4.**
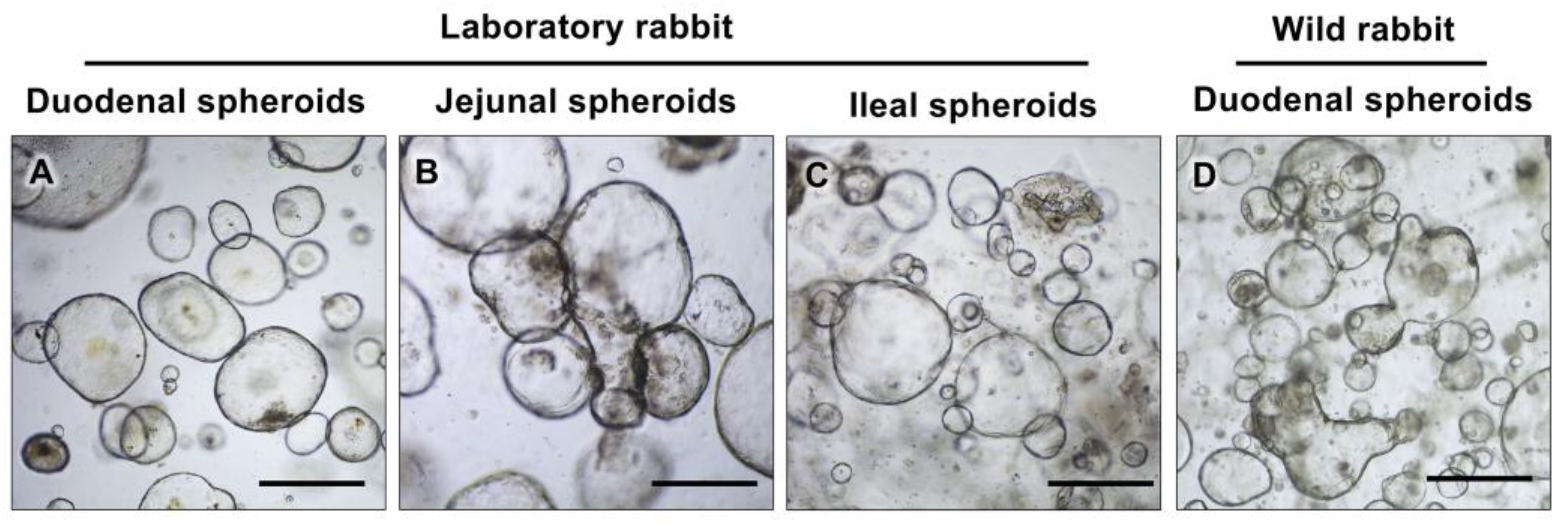
Spheroid morphology. Rabbit intestinal spheroids from different tissue segments have similar morphology. L-WRN-conditioned medium supported the growth of spheroids from various segments of small intestines, (A) duodenum, (B) jejunum and (C) ileum from a laboratory rabbit, and (D) duodenum from a wild rabbit. Scale bars = 500 μm.

**Fig 5.**
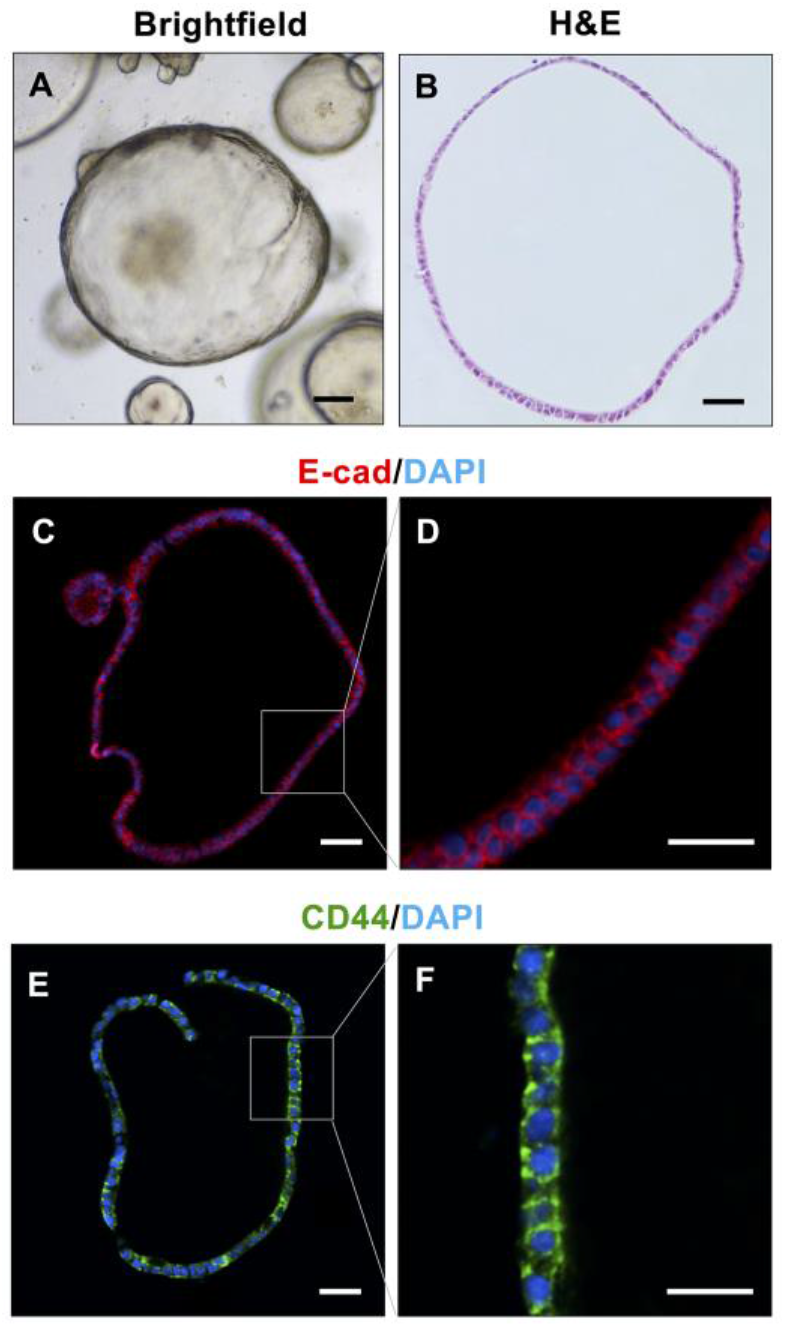
Identification of stem cells in rabbit duodenal spheroids. Rabbit duodenal spheroids visualised with (A) brightfield and (B) H&E staining. Spheroids were immuno-stained with (C, D) an epithelial cell marker, E-cadherin (red), and (E, F) a stem cell marker, CD44 (green). Nuclei were counterstained with DAPI (blue). Scale bars = 100 μm.

While establishing the most suitable passaging conditions for rabbit spheroids, we were surprised to find that different methods of cell dissociation lead to differences in spheroid morphology and viability. The mechanical shearing of spheroids using hypodermic needles resulted in spheroid fragments that spontaneously differentiated into ‘multilobular organoids’ that contained goblet cells and survived for only 8 days in proliferation medium (data not shown). In contrast, enzymatic dissociation with the TrypLE™ Express enzyme produced ‘regular’ spheroids that could be sub-cultured at least 17 times.

### ROCK and TGF-β inhibitors are important for growing rabbit intestinal spheroids

Miyoshi and Stappenbeck demonstrated that although both ROCK and TGF-β inhibitors are required in early passages of both human and mouse spheroid cultures, these inhibitors were no longer required in later passages [27]. However, we found that rabbit spheroid cultures grew best in the continued presence of both ROCK and TGF-β inhibitors (as quantified by measuring the diameter of spheroids; Fig 6A–E). Without any inhibitor, or with the addition of only the ROCK or TGF-β inhibitor, spheroids grew to a similar size (58.48 ±28.81, 52.92 ± 30.57 and 51.54 ±27.17 μm, respectively; Fig 6A–C). However, in the presence of both ROCK and TGF-β inhibitors, the spheroids grew to a significantly larger size (172.85 ± 101.54 μm; *p* < 0.05; Fig 6D, E). These data suggest a synergistic effect of ROCK and TGF-β inhibitors.

**Fig 6.**
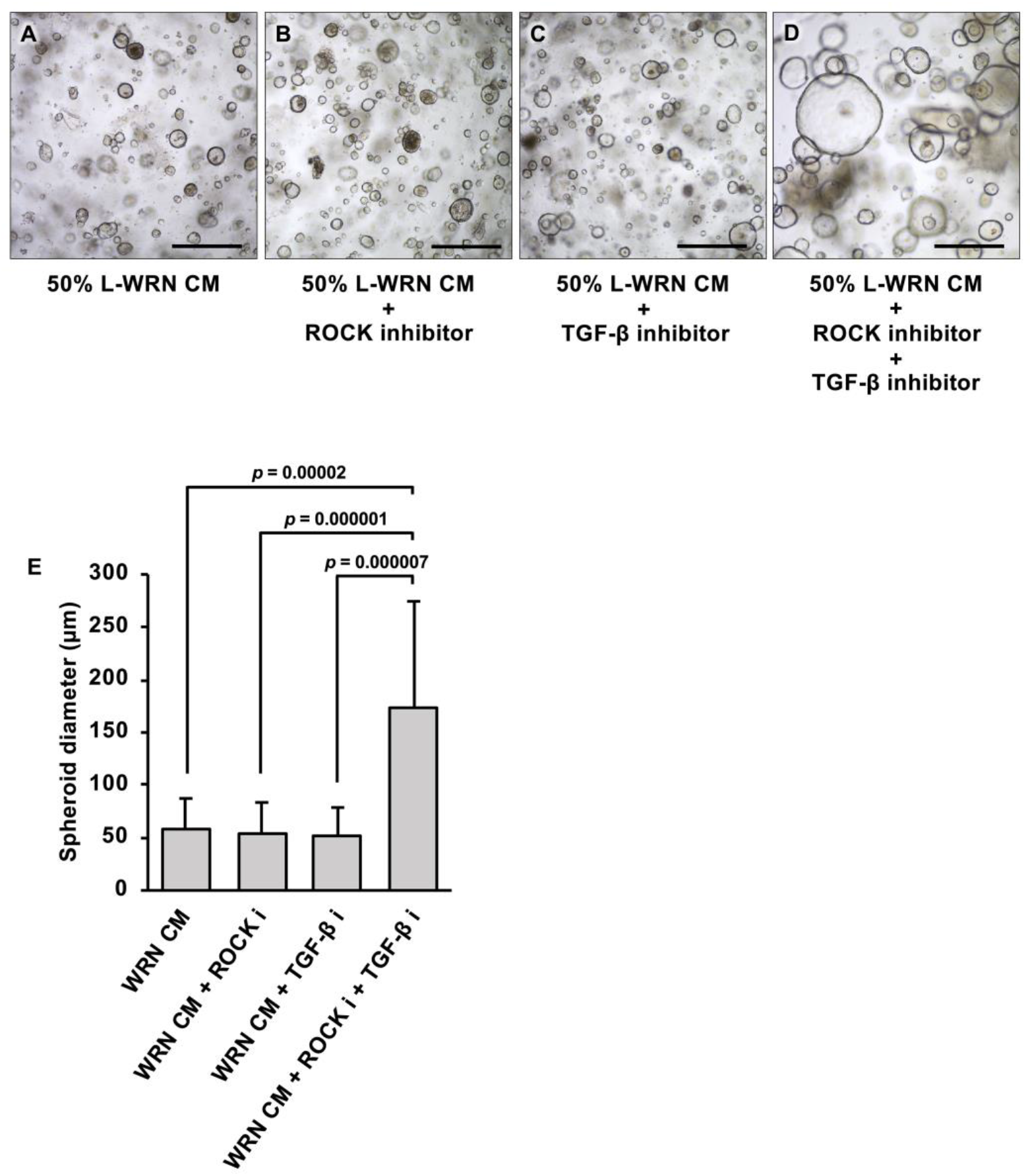
Effect of ROCK and TGB-β inhibitors on the growth of rabbit intestinal spheroids. Different combinations of proliferation medium were tested to optimise cell culture conditions for spheroid formation. Spheroids were passaged and cultured in (A) L-WRN-conditioned medium without any additional inhibitors, (B) L-WRN-conditioned medium with ROCK inhibitor, (C) L-WRN-conditioned medium with TGB-β inhibitor and (D) L-WRN-conditioned medium with ROCK and TGB-β inhibitors. Scale bars = 500 μm. (E) The diameter of rabbit intestinal spheroids was measured after 5 days in culture and compared between treatment groups. The height of the columns represents the average diameter of 50 spheroids from three wells (error bars indicate the standard error of the mean). Student’s t-test was performed to determine the significance of the observed size differences and *p* values are given for all significant (*p* < 0.05) differences.

### Differentiation of rabbit duodenal spheroids to organoids

In this study, our differentiation medium consisted of diluted L-WRN-conditioned medium, a Notch signalling inhibitor (DAPT) and a ROCK inhibitor. Spheroid cultures were grown for 3–4 days before the proliferation medium was exchanged for differentiation medium. Differentiation medium differed from proliferation medium by the reduction of WRN stem cell growth factors, the addition of DAPT, a Notch signalling inhibitor, and the removal of the TGF-β inhibitor. Four days after initiation of the differentiation process, duodenal spheroids developed cellular characteristics that resembled an intestinal epithelium (Fig 7A, 8A, B). We detected protrusions that appeared at the periphery of spheroids, which denotes intestinal organoid maturation. In previous studies using human small intestinal organoids, two distinct morphologies were identified; cystic (sometimes also referred to as enterospheric [32]) and multilobular [33,34]. Multilobular organoids have one or multiple buds, whereas those without crypt-like protrusions are referred to as cystic. We found both morphologies in our rabbit duodenal organoid cultures (Fig 7); an examination of 136 organoids revealed that 85% were cystic (Fig 7A) and 15% were multilobular (Fig 7B).

**Fig 7.**
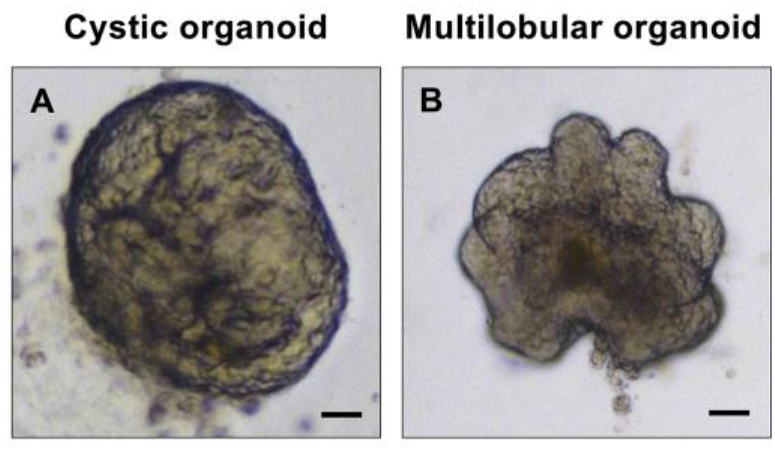
Morphology of differentiated rabbit duodenal organoids. Organoids that formed after four days in differentiation medium were observed to have either a (A) cystic or (B) multilobular morphology.

**Fig 8.**
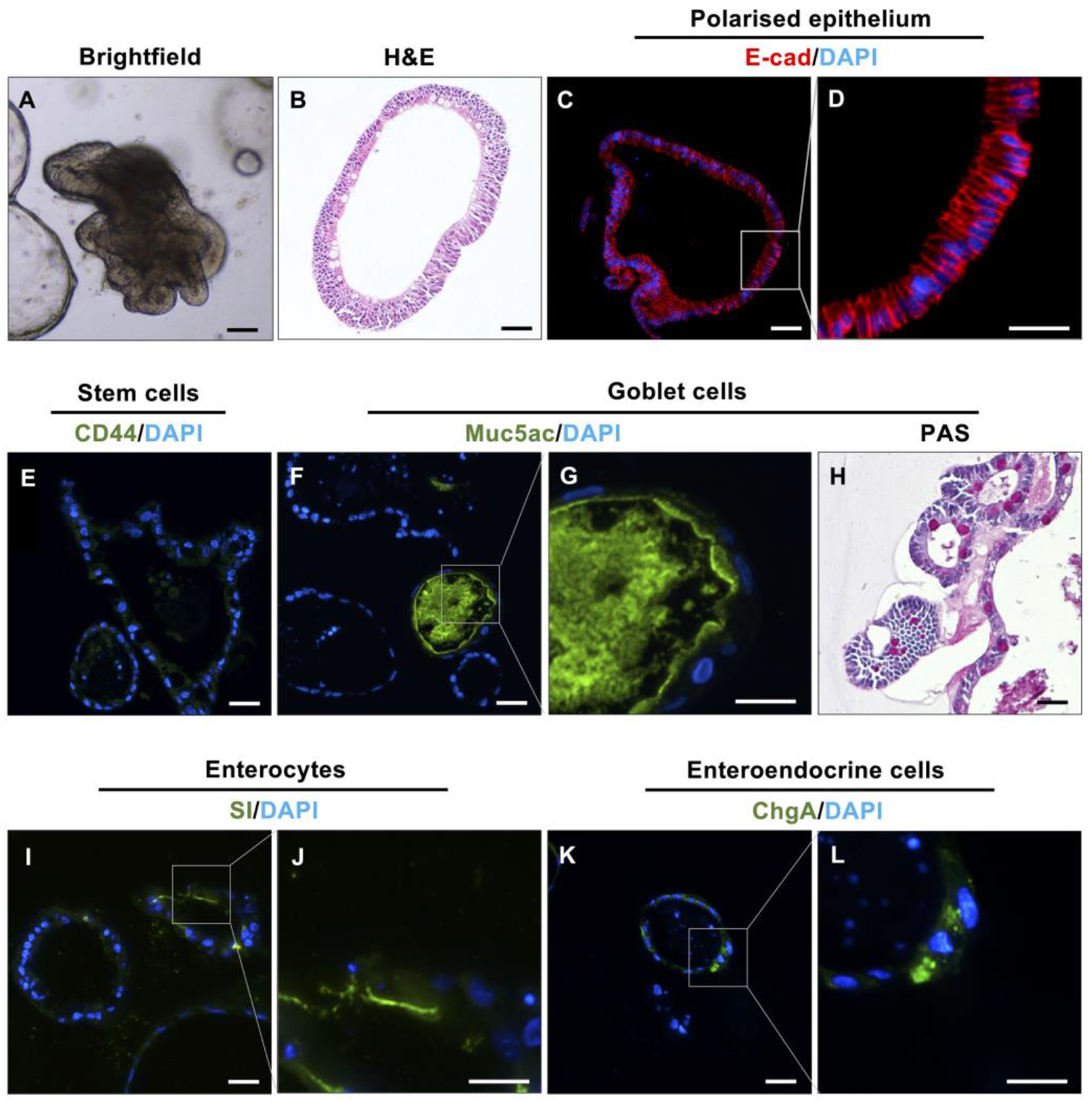
Identification of different cell types in rabbit duodenal organoids. Visualisation of rabbit duodenal organoids in (A) brightfield and (B) H&E staining. Immunofluorescence staining indicated multiple different cell types within the intestinal epithelium after four days in differentiation medium. (C, D) E-cadherin staining (red) shows the presence of polarised epithelial cells. (E) The absence of the stem cell marker CD44 suggests that all stem cells differentiated. (F, G) Muc5ac (green) and (H) PAS staining (magenta) indicates mucus production by goblet cells. (I, J) sucrase-isomaltase (SI) staining (green) visualises the brush border of mature enterocytes. (K, L) Chromogranin A (ChgA) staining (green) demonstrates the presence of enteroendocrine cells. Nuclei were counterstained with DAPI (blue). Scale bars = 100 μm (A–D) and 200 μm (E–L).

Based on immunofluorescence staining, mature rabbit duodenal organoids showed the typical hallmarks of differentiated intestinal epithelial lineages. E-cadherin staining demonstrated a thicker epithelium lining compared to spheroids and polarised columnar epithelium (Fig 8C). We could no longer detect the stem cell population (i.e., CD44^+^) that was previously present in the spheroids (Fig 8D). The stem cells differentiated into several intestinal epithelial cell types, including goblet cells (PAS^+^, Muc5ac^+^; Fig 8E, F), enterocytes (SI^+^; Fig 8G) and enteroendocrine cells (ChgA^+^; Fig 8H).

To complement our protein expression analysis, we used RT-qPCR to assess mRNA expression of genes related to intestinal maturity and differentiation. Rabbit intestinal organoids grown in differentiation medium exhibited a decrease in intestinal stem cell-associated transcripts, including *AXIN2* (Wnt signalling activity) and *LGR5* (intestinal stem cells) (Fig 9). In contrast, the expression of genes associated with mature intestinal epithelial cells, such as *MUC5AC, SI, CHGA* and *LYZ* were highly upregulated in differentiated organoid culture (Fig 9).

**Fig 9.**
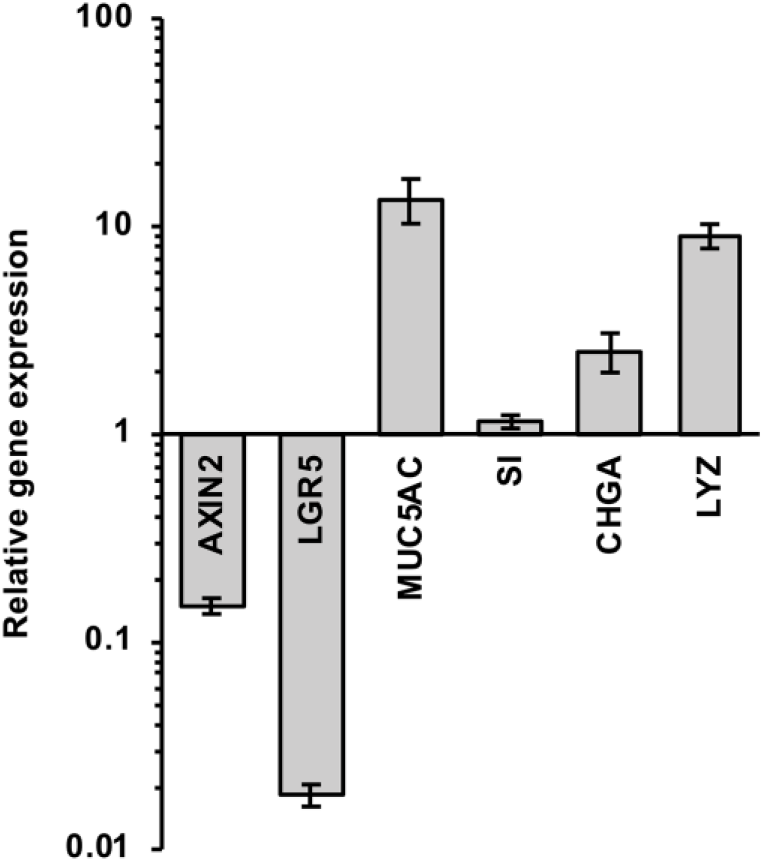
Gene expression in differentiated rabbit duodenal organoids relative to undifferentiated rabbit duodenal spheroids. RT-qPCR analysis was performed to compare mRNA levels of selected stem cell-associated (*AXIN2, LGR5*) and maturation-associated genes (*MUC5A, SI, CHGA, LYZ*) between differentiated and undifferentiated spheroids and organoids. Data are presented as fold change (2^-ΔΔCt^) between undifferentiated spheroids versus differentiated organoids calculated from three independent experiments with three technical replicates for each assay. Columns heights and error bars represent the mean fold change in expression levels and standard error of the mean, respectively.

### Rabbit duodenal spheroid-derived monolayer cultures contain differentiated cells

To establish monolayer cultures, rabbit duodenal spheroids were digested with TrypLE™ Express enzyme and the dissociated cells were plated onto coated culture plates. We found that cells adhered best to surfaces coated with 100 μg/ml of collagen type IV; the use of a lower collagen concentration resulted in poor cellular attachment. Furthermore, we found that a seeding density of approximately 7 *×* 10^4^ cells/well was optimal for starting rabbit duodenal monolayer cultures in both 96-well plates or Transwell^®^ membrane inserts.

Immunofluorescence staining of monolayer cells grown in differentiation media for four days could no longer detect the expression of the stem cell marker CD44 (Figs 8E and 10A) but revealed the presence of the same differentiated intestinal epithelial cell lineages that were found in the organoids (Fig 10B–E). Muc5ac (Fig 10B) and PAS staining (Fig 10C) demonstrated the presence of goblet cells, whereas SI (Fig 10D) and ChgA (Fig 10E) indicated the presence of enterocytes and enteroendocrine cells, respectively. To corroborate these findings, we analysed the expression of selected genes by RT-qPCR (Fig 11). We found reduced expression levels for the stem-cell-associated transcripts *AXIN2* and *LGR5*, and increased levels for *MUC5AC, SI* and *CHGA*, which suggests that stem cells differentiated into goblet cells, enterocytes and enteroendocrine cells. However, we did not find an increase in the expression of *LYZ*, suggesting that our monolayer cultures did not contain Paneth cells.

**Fig 10.**
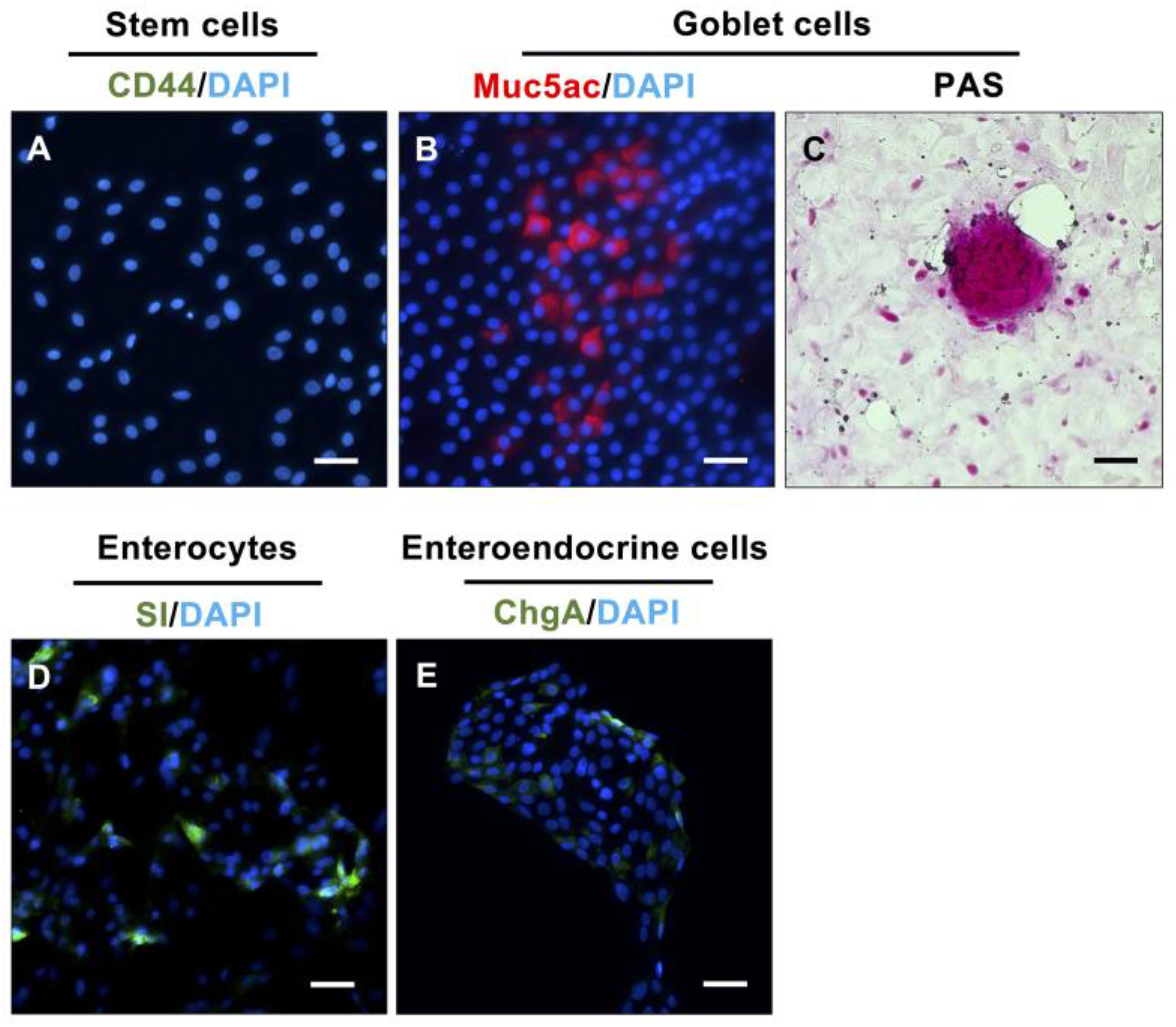
Identification of different cell types in rabbit duodenal monolayer cultures. Immunofluorescence staining showed cell differentiation after four days in differentiation medium. (A) Monolayer cultures showed no expression of the stem cell marker CD44. (B–E) Muc5ac (red), PAS (magenta), SI (green) and ChgA (green) indicates the presence of goblet cells, enterocytes and enteroendocrine cells, respectively. Nuclei were counterstained with DAPI (blue). Scale bars = 100 μm.

**Fig 11.**
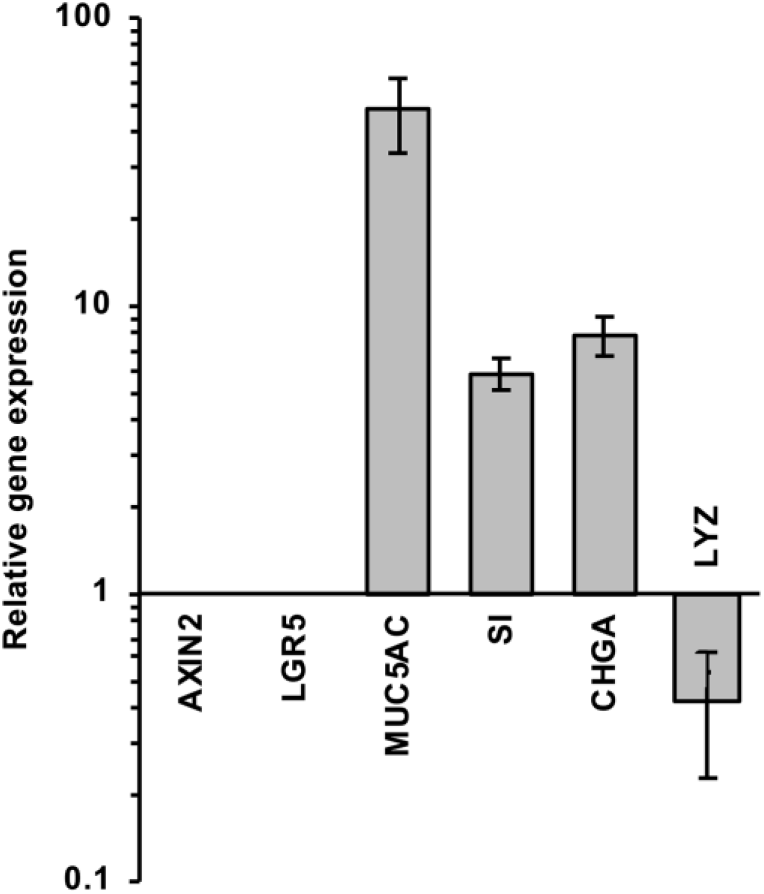
Gene expression in undifferentiated and differentiated rabbit duodenal cell monolayers. Cell monolayers were grown either in the presence of proliferation medium (‘undifferentiated cultures’) or differentiation medium (‘differentiated cultures’). RT-qPCR analysis was performed to analyse mRNA levels of stem cell-associated (*AXIN2, LGR5*) and differentiation-associated genes (*MUC5A, SI, CHGA, LYZ*) between undifferentiated and differentiated cultures. The data were calculated from three independent experiments with three technical replicates. Columns heights and error bars indicate the fold change (2^-ΔΔCt^) in expression levels and standard errors of the mean.

## Discussion

Here, we report robust methods to generate rabbit intestinal organoids and monolayer cultures. This includes protocols for the generation, propagation and long-term cryogenic storage of spheroids from a variety of small intestinal tissues, including the duodenal, jejunal, and ileal segments. We were able to culture spheroids for at least 17 passages without any notable changes in morphology or growth rate. We were also able to freeze and revive the frozen spheroids with minimal loss of cell viability. Our newly established rabbit organoid cultures reproduce many characteristics of the parental tissue. For example, we detected (i) the presence of multiple mature cell types such as enterocytes, enteroendocrine cells, goblet cells and Paneth cells, (ii) brush borders on enterocytes, (iii) the production of mucin by goblet cells and (iv) the synthesis of lysozyme by Paneth cells. Our observations are in line with previous reports on intestinal organoids that were derived from other species, including mice [35] and pigs [36].

We found that rabbit duodenal organoids showed both cystic and multilobular morphologies, but most organoids were cystic. Both morphologies have previously been reported for human intestinal organoids [33]. The mechanism that drives multilobular organoid morphology remains unknown, but Paneth cells are suspected to regulate and determine the fate of stem cell differentiation during organogenesis or upon intestinal damage through the secretion of Wnt3 [17]. In cell culture, the addition of Wnt growth factor allows stem cells to create their own niche, self-organise and form organoids. Interestingly, it has been reported that a knock down of the *Wnt3* gene resulted in murine intestinal organoids that had a less pronounced multilobular morphology [17]. It would be interesting to determine if enhanced Wnt signalling, e.g., through higher Wnt concentrations or the use of a homologous (rabbit) cytokine would result in a higher proportion of multilobular rabbit intestinal organoids.

The growth factors Wnt, R-spondin and Noggin are the three main growth factors required to maintain a stem cell population in mouse or human spheroid cultures [27]. In several previous studies, L-WRN-conditioned media that were supplemented with ROCK and a TGF-β inhibitor were used to generate intestinal spheroids from human, mouse, cow, cat and dog cells [27,30,37,38]. Therefore, we expected that a similar L-WRN-conditioned medium would also be essential for the establishment of intestinal spheroid cultures from rabbits. While testing different culture media, we found that adding both ROCK and TGF-β inhibitors significantly improved the growth of spheroids from the small intestine of rabbits. Thus, our culture conditions are somewhat different to what has previously been used for establishing small intestinal spheroids from mice [27]. The ROCK inhibitor that was used in this study, Y-27632, inhibits the two isoforms of the Rho-associated coiled-coil-containing protein kinase ‘ROCK’, i.e., ROCK 1 and ROCK 2. Several ROCK substrates are involved in the execution and possibly also in the initiation of apoptosis. Y-27632 has been used to control stress conditions and enhance cell recovery in primary cell isolation and cryopreservation [39]; the inhibitor has also been used to reduce dissociation-induced apoptosis in human embryonic stem cell culture [39,40] and primary primate corneal endothelial cells [41]. However, the response to Y-27632 is cell type-specific and depends on the apoptotic stimulus, which may explain why using Y-27632 is beneficial in some spheroid cultures, including ours, but not in others.

SB431542 is a selective inhibitor that blocks the activity of the TGF-β type I receptor-like kinases ALK4, ALK5 and ALK7 [42]. The BMP/TGF-β signalling pathway is responsible for intestinal epithelial cell differentiation [43]. The addition of TGF-β antagonist such as SB431542 in culture medium prevented spontaneous differentiation and maintained the stemness of mouse embryonic stem cells [44]. Our results show that blocking both ROCK and TGF-β signals is required for the prolonged propagation of rabbit intestinal spheroids.

As WRN growth factors promote stem cells in spheroid cultures, these factors are no longer required for the cultivation of differentiation organoids. Consequently, the removal of these factors resulted in stem cell differentiation. Differentiation medium supplemented with DAPT, a Notch signalling inhibitor, stopped stem cells from proliferating and promoted human intestinal organoid differentiation [37,45]. These same conditions were found to facilitate differentiation in our rabbit duodenal organoid cultures.

Rabbit intestinal organoids may be utilised to facilitate functional studies of important human gastrointestinal tract diseases. The rabbit, for example, is an important model for inflammatory bowel diseases, such as ulcerative colitis. Rabbits show a similar colonic response to that of humans when exposed to inflammatory agents [46]. Thus, rabbit intestinal organoids will be a useful tool to complement animal studies on inflammatory bowel disease. Other important human gastrointestinal tract diseases are caused by bacteria (e.g., *Shigella, Helicobacter* and *Salmonella*) and viruses (e.g., noroviruses and rotaviruses). Infectious disease models for these pathogens traditionally rely on mice and rats, but rabbits have also been used. For example, rabbits are used as a model to study human bacillary dysentery or shigellosis, a colonic and rectal infection caused by *Shigella* spp. Infecting young rabbits leads to bloody diarrhea [47,48], one of the most important features of human shigellosis that cannot be reproduced in other animal models [49,50]. The availability of rabbit organoids will allow more detailed studies of this disease.

The generation of rabbit small intestinal organoids and intestinal monolayer cultures derived from spheroids will facilitate more detailed investigations into the physiology of the lagomorph gastrointestinal tract. Rabbits and other lagomorphs have evolved a digestive system that is radically different to that of other, better known herbivores [51]. The underlying mechanisms for their evolutionary success are not fully understood and organoids will aid further insight. Organoid culture systems not only provide a more biologically relevant model system compared to traditional 2D cell cultures, but they will also reduce the use of animals for research purposes.

## Acknowledgements

We thank the members of rabbit biocontrol team (Elena Smertina, Nina Huang, Maria Jenckel, Tegan King, Ina Smith, Roslyn Mourant and Melissa Piper), Marcin Büler from CSIRO Land and Water and members of the Estes’ laboratory at the Baylor College of Medicine (Umesh Karandikar, Khalil Ettayebi, Victoria Tenge and Shih-Ching Lin) for experimental assistance and advice. Finally, we thank Sarron Randall-Demllo and Peter Kerr from CSIRO Health and Biosecurity and Kerry Mills for proofreading the manuscript. This project was funded by a CSIRO OCE Postdoctoral Fellowship (2017-18, Round 1) with additional support from the Mitigating Invasive Species and Diseases program of CSIRO Health and Biosecurity.

